# Correction of β3 integrin haplo-insufficiency by CRISPRa normalizes cortical network activity

**DOI:** 10.1101/664706

**Authors:** Fanny Jaudon, Agnes Thalhammer, Lorenzo A. Cingolani

**Affiliations:** Center for Synaptic Neuroscience and Technology (NSYN), Istituto Italiano di Tecnologia (IIT), Genoa, Italy; IRCCS Ospedale Policlinico San Martino, Genoa, Italy; Department of Life Sciences, University of Trieste, Trieste, Italy

## Abstract

The extracellular matrix (ECM) plays an essential role in regulating the function of neuronal networks. In many cell types, the ECM exerts its effects through the transmembrane receptors integrins. Here, we investigate whether neuronal integrins regulate network excitability. Specifically, we focus on β3 integrin, which has been associated to autism spectrum disorder and whose expression in neurons is regulated by activity. We have designed CRISPRa tools to titrate β3 integrin expression in neurons. By using multi-electrode arrays and Ca^2+^imaging, we show that β3 integrin boosts network excitability and synchrony in primary cortical neurons. Crucially, CRISPRa could compensate precisely for β3 integrin haplo-insufficiency, thereby rebalancing level and correlation of neuronal activity. By contrast, rescue strategies based on exogenous gene expression resulted in hyper-active networks firing in unison. Thus, regulation of β3 integrin by CRISPRa provides a precise and efficient system for modulating neuronal network function.

## INTRODUCTION

Growing evidence indicates that integrins play an essential role in regulating synaptic connectivity and plasticity in response to extracellular cues (Ferrer-Ferrer and Dityatev, 2018; Kerrisk et al., 2014; Lilja and Ivaska, 2018; Park and Goda, 2016; Thalhammer and Cingolani, 2014). β1 and β3 integrins, the two best studied neuronal integrins, have divergent functions in the brain. While β1 integrin is implicated in basal synaptic transmission and LTP stabilization (Babayan et al., 2012; Chan et al., 2006; Huang et al., 2006; Liu et al., 2016; Warren et al., 2012), β3 integrin is required for homeostatic synaptic plasticity, which maintains network activity within an optimal physiological range. Specifically, activity deprivation, to induce homeostatic upscaling of excitatory synaptic currents, up-regulates also β3 integrin surface expression, while genetic ablation of this integrin prevents selectively homeostatic upscaling (Cingolani and Goda, 2008; Cingolani et al., 2008; McGeachie et al., 2012). Thus, some aspects of activity-dependent adjustment of neuronal network activity involve integrin signaling.

It is however not known whether changes in integrin expression *per se* affect network excitability. Notably, variations in expression and function of β3 integrin have been associated to autism spectrum disorder (ASD) (Cantor et al., 2005; Dohn et al., 2017; O’Roak et al., 2012; Weiss et al., 2006) and deficits in homeostatic synaptic plasticity may underlie some aspects of ASD (Bourgeron, 2015; Nelson and Valakh, 2015). It would therefore be greatly beneficial for investigating the physiological and pathological roles of integrins in the brain to develop effective strategies for fine-tuning their neuronal expression.

The CRISPR/Cas9 technology, originally used for genome editing, has recently been repurposed for regulating gene expression. A nuclease deficient Cas9 (dCas9), fused to transcription activators, such as VP64, can be directed via guide RNAs (gRNAs) to the promoter region of a gene of interest to enhance its transcription, an approach known as CRISPR activation (CRISPRa; **Fig S4C**; (Chavez et al., 2015; Konermann et al., 2015; Tanenbaum et al., 2014). Despite the potential of CRISPRa, its use in neurons remains however challenging (Savell et al., 2019; Zhou et al., 2018).

Using multi-electrode arrays (MEAs) and Ca^2+^ imaging in primary cortical neurons from WT, *Itgb3* heterozygous (Het) and KO mice, we show that β3 integrin boosts cortical network activity and synchrony. Further, we implement CRISPRa to compensate precisely for β3 integrin haplo-insufficiency, thereby rebalancing level and correlation of neuronal activity. By contrast, classical rescue strategies based on the expression of exogenous β3 integrin resulted in hyper-active networks firing in unison. Thus, CRISPRa provides a flexible system for fine-tuning expression of endogenous genes within a physiological range and modulation of neuronal network activity.

## RESULTS

### Reducing β3 integrin expression lowers neuronal activity

To address whether β3 integrin regulates network activity, we used primary cortical cultures derived from *Itgb3*^*+/*+^ (WT), *Itgb3*^*+/*−^ (Het) and *Itgb3*^−*/*−^ (KO) mice. RT-qPCR, Western blots and confocal imaging showed that both mRNA and protein levels for β3 integrin were reduced by ∼50% in Het, as compared to WT, and were undetectable in KO (**Fig S1A-D**). We next asked whether these differences affected cortical network activity. To this end, we used a red-shifted genetically encoded Ca^2+^ indicator, jRCaMP1b (Dana et al., 2016), to monitor spontaneous fluorescence signals in neurons (**Fig 1**), which can be interpreted as correlates of neuronal activity (Inoue et al., 2019). Somatic jRCaMP1b transients had a favorable signal-to-noise ratio (**Fig 1A, B**). While we did not detect significant differences in the size of the spontaneous fluorescence responses between the three genotypes, we observed a ∼38% reduction in frequency in both Het and KO neurons (**Fig 1C**).

**Fig 1.**
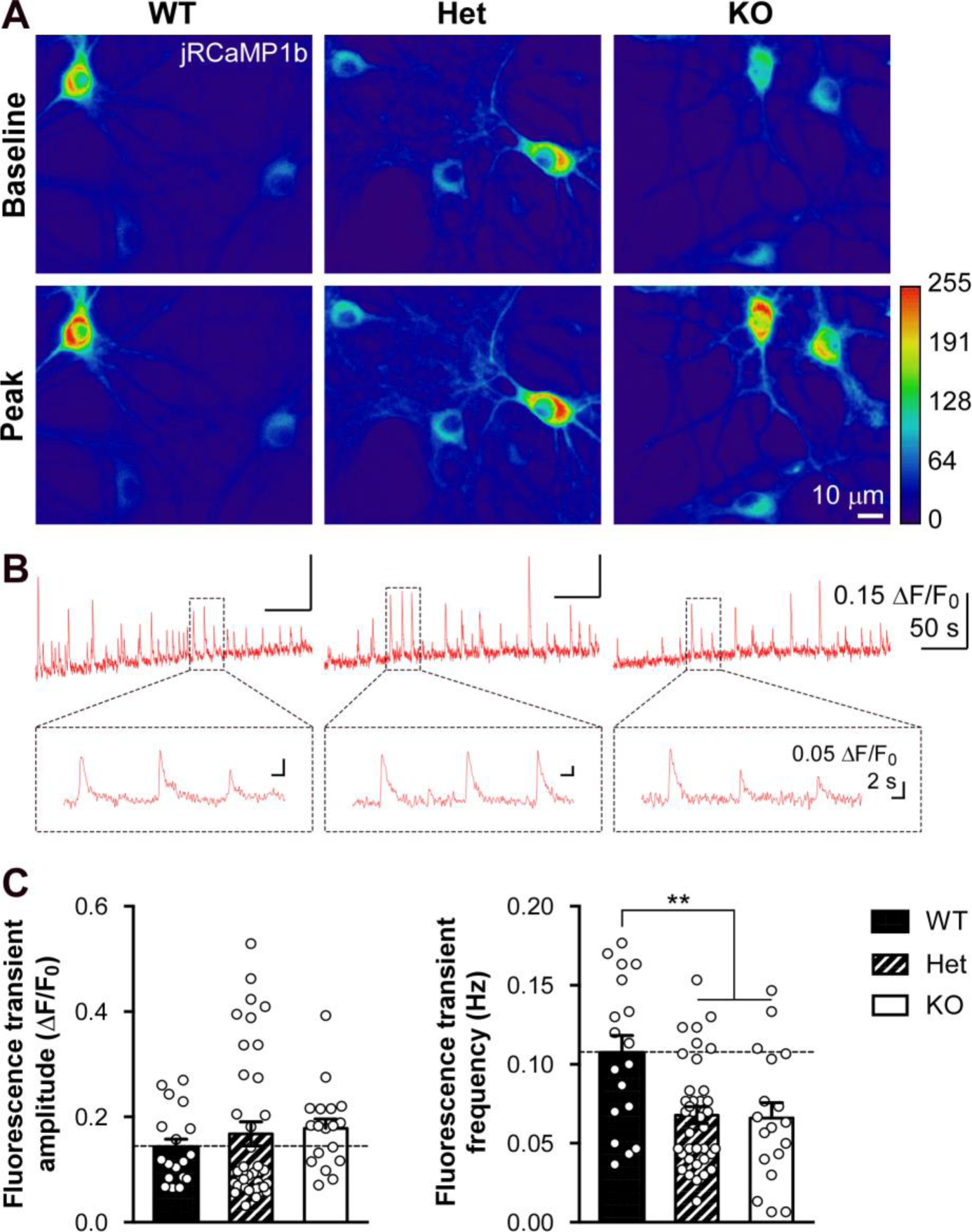
Spontaneous activity is reduced in β3 integrin Het and KO cortical neurons. **(A)** Representative jRCaMP1b fluorescence transients in response to spontaneous network activity in primary cortical cultures at 16 DIV. Images are average of six consecutive frames during baseline and at the peak of the largest transient. **(B)** Spontaneous somatic jRCaMP1b responses over 5 min. Inset, higher magnification, showing good signal-to-noise ratio. **(C)** Quantification of experiments as in (A-B) for amplitude and frequency. Cortical neurons from *Itgb3* Het and KO mice exhibit lower frequency of spontaneous fluorescence transients (**p<0.01, one-way ANOVA followed by Tukey post-test, n=18, 38 and 18 recordings for WT, Het and KO, respectively; 7-8 independent cultures). Data are presented as mean±SEM; dots represent individual recordings. See also Fig S1.

Thus, a decrease in β3 integrin expression by 50%, as observed in Het neurons, is as effective as a complete ablation of this protein in reducing network activity.

### CRISPRa allows for fine-tuning of β3 integrin expression in neurons

We next attempted to enhance transcription of β3 integrin as a means to regulate neuronal activity. To this end, we used CRISPRa where dCas9 is fused to the transcriptional activator VP64 (Perez-Pinera et al., 2013). We designed three gRNAs targeting the *Itgb3* promoter and expressed them together with dCas9-VP64 and EGFP (**Fig S1F**) in the murine N2a cell line (**Fig S1G**). As quantified by RT-qPCR, all gRNAs induced a significant 3 to 4-fold increase in β3 integrin expression, with gRNA 3 being the most effective (**Fig S1H**). While no additive effect was observed when combining gRNAs 3 and 1, likely because of the steric hindrance between the two target sequences (**Fig S1F**), co-expression of gRNAs 2 and 3 increased β3 integrin expression by 8-fold (**Fig S1H**).

As functional readout for the detected differences in mRNA expression, we performed a cell attachment assay. CRISPRa significantly increased adhesion of N2a cells to the β3 integrin-specific ligand fibronectin (**Fig S1I, J**).

To address whether CRISPRa can be used to regulate β3 integrin expression in neurons, we infected primary cortical cultures from WT, *Itgb3* Het and KO mice with lentiviruses expressing a gRNA together with dCas9-VP64 and EGFP (**Fig 2A**). gRNA 3 induced a 2 to 2.5-fold increase in β3 integrin mRNA in both WT and Het, but not in KO, where both copies of the target gene are missing. Hence, CRISPRa allows for β3 integrin haplo-insufficiency in Het to be nearly perfectly recovered to WT values (**Fig 2B**).

**Fig 2.**
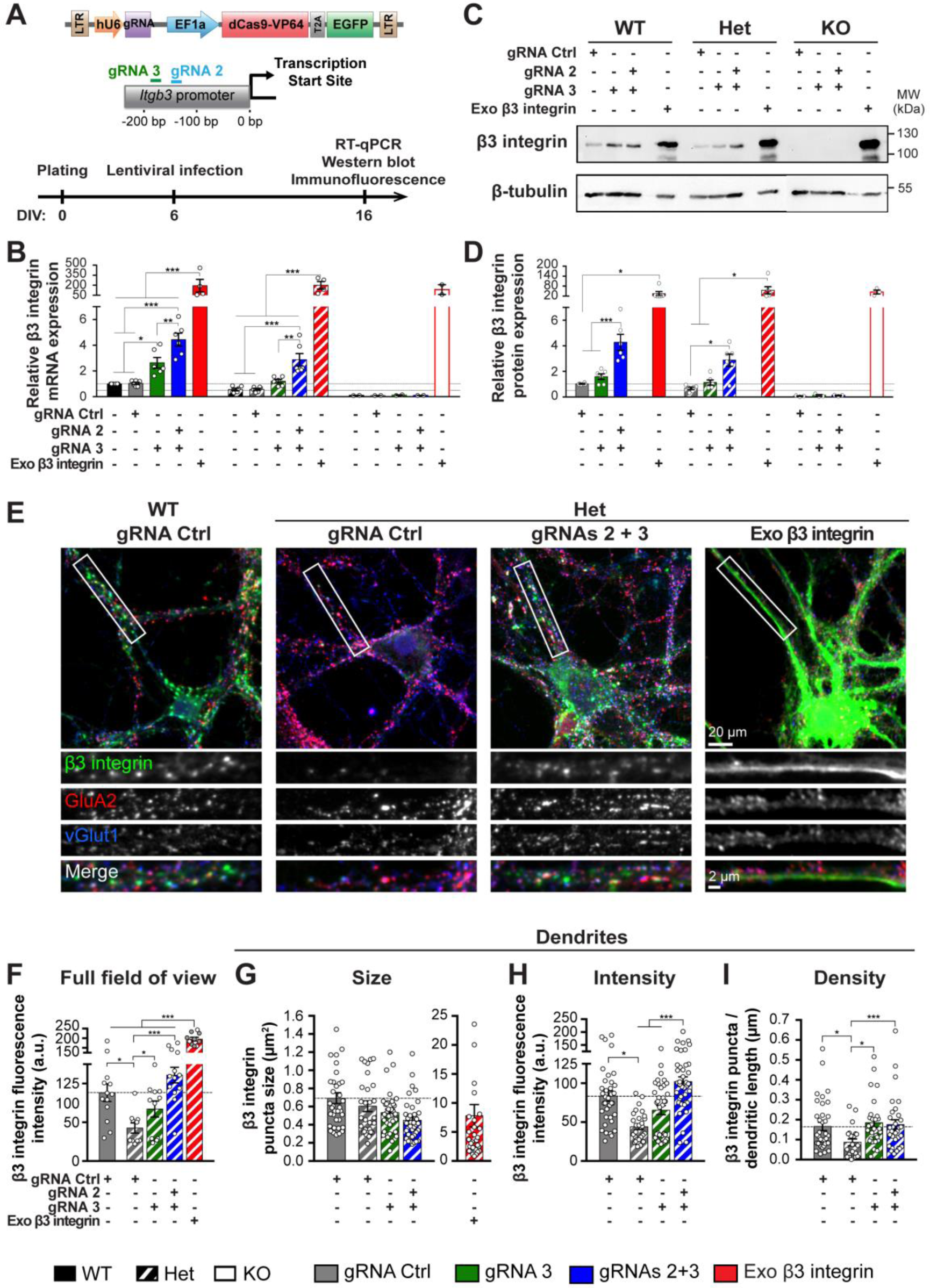
Normalization of β3 integrin expression by CRISPRa. **(A)** Lentiviral construct, scheme of the gRNA targets on the *Itgb3* promoter and experimental timeline. **(B)** RT-qPCR quantification of β3 integrin mRNA expression in WT, *Itgb3* Het and KO cortical neurons transduced with the indicated constructs. (n=6, 6 and 2 independent cultures for WT, Het and KO, respectively; 2 technical replicates each). **(C)** Representative Western blots of membrane fractions. **(D)** Quantification of experiments as in (C) (n=6, 6 and 3 independent cultures for WT, Het and KO, respectively; 2 technical replicates each). **(E)** Representative confocal images of primary cortical neurons from WT and Het cultures expressing the indicated constructs. β3 integrin, the glutamate receptor subunit GluA2 and the presynaptic marker vGlut1 are shown in false colors; infection was confirmed by EGFP. **(F)** Quantification of β3 integrin fluorescence intensity for the full field of view for experiments as in (E), indicating that CRISPRa elevates β3 integrin expression in Het to WT values while exogenous expression of β3 integrin increases several-fold the signal for this protein (the fluorescence intensity for Het+exogenous β3 integrin is a lower estimate because of pixel saturation; gray filled circles indicate pixel saturation >30%). **(G)** Dendritic puncta size for β3 integrin; expression of exogenous β3 integrin resulted in large dendritic areas of saturated signal, which were no further analyzed. **(H-I)** Quantification of the effects of the indicated gRNAs on dendritic puncta intensity (H) and number (I; n=13, 10, 11, 13 and 10 field of views from 3 independent cultures each for WT+gRNA Ctrl, Het+gRNA Ctrl, Het+gRNA 3, Het+gRNAs 2+3 and Het+exogenous β3 integrin, respectively) Data are presented as mean±SEM; dots represent individual values (*p<0.05, **p<0.01, ***p<0.001, one-way ANOVA followed by Tukey post-test). See also Fig S1, S2 and S3.

Like in N2a cells, combining gRNAs 3 with 2 led to a stronger increase in β3 integrin mRNA in both WT and Het (3 to 4.5-fold, **Fig 2B**). We next compared CRISPRa with a classical lentiviral overexpression approach (McGeachie et al., 2012), and found that the amount of exogenous β3 integrin mRNA was ∼200-fold higher than endogenous WT values.

Changes in mRNA were paralleled at the protein level. As assessed by Western blots, CRISPRa restored the amount of WT protein in Het neurons, whereas expression of exogenous β3 integrin induced a 40 to 50-fold increase in protein expression (**Fig 2C, D**). Unlike overexpression, CRISPRa allows therefore for fine-tuning of neuronal β3 integrin levels within a physiological range.

Cas9 has been shown to bind to off-targets but whether this leads to off-target effects remains unclear (Kuscu et al., 2014; Wu et al., 2014). Although none of the top predicted off-targets for gRNAs 2 and 3 were located on a promoter region (**Fig S2A**), we checked whether the two gRNAs were prone to off-target effects by evaluating the binding of dCas9 to the top ten off-targets for each gRNA using chromatin immunoprecipitation followed by qPCR (ChIP-qPCR; **Fig S2B-C**). Relative to gRNA Ctrl, gRNAs 2 and 3 induced a 25-fold enrichment in dCas9 binding at the *Itgb3* promoter (**Fig S2C**), confirming dCas9 binding to the target site. A small but significant enrichment in dCas9 binding was detected for two predicted off-target sites (#9 of gRNA 2 and #6 of gRNA 3; **Fig S2C**). This was however without effect on the expression of the genes at these loci (*Msra* and *Braf*; **Fig S2D**).

### Normalization of dendritic expression of β3 integrin by CRISPRa

β3 integrin is expressed in dendrites in tight apposition to synaptic markers (Cingolani et al., 2008; Pozo et al., 2012). We therefore assessed, by confocal microscopy, the localization of β3 integrin upon CRISPRa enhancement of transcription. The reduced expression of β3 integrin in Het neurons could be augmented to WT levels by CRISPRa in both soma and dendrites (**Fig 2E, F** and **H**) whereas no effect was observed in KO neurons (**Fig S1E**). The size and density of β3 integrin clusters along dendrites was largely preserved (**Fig 2E, G** and **I**). By contrast, expression of exogenous β3 integrin led to widespread and ectopic localization of the protein (**Fig 2E-G**). None of the experimental conditions affected expression of and co-localization with the synaptic markers GluA2 and vGlut1 (**Fig S3A-H**).

In summary, β3 integrin maintains its expression pattern when the activity of its endogenous promoter is enhanced by CRISPRa, thereby precisely reversing haplo-insufficiency of this gene.

### Normalization of cortical network activity by CRISPRa

We next asked whether enhancing expression of β3 integrin by CRISPRa regulates cortical network activity. To this end, we recorded the activity of high density cultures grown on MEAs. For all experimental conditions, spontaneous basal activity was characterized by random spikes and bursts (**Fig 3A, B**). Under control conditions, the overall activity of the network was lower for Het and KO cultures than for WT cultures (**Fig 3A-G**), thus corroborating the jRCaMP1b results (**Fig 1**).

**Fig 3.**
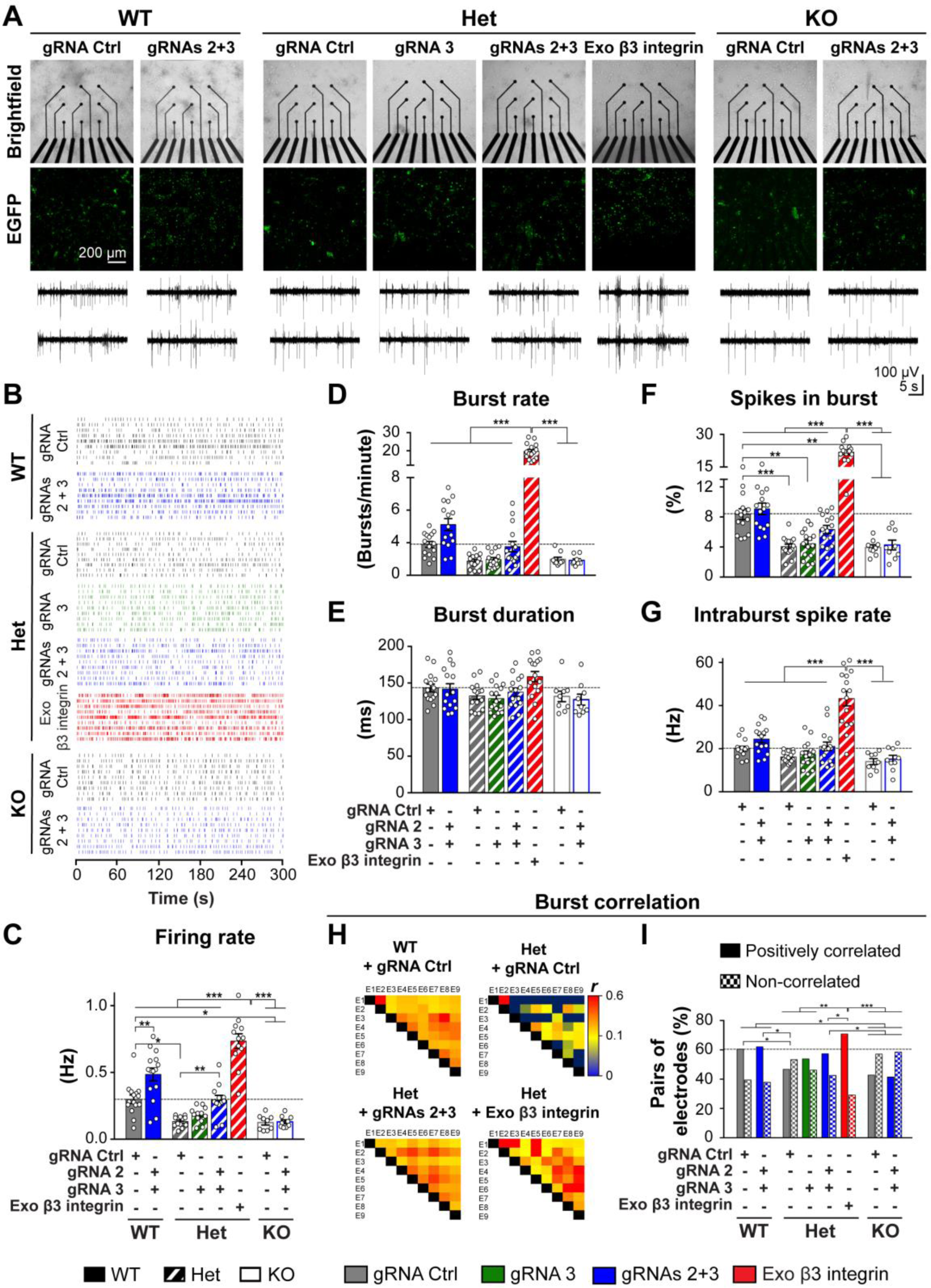
CRISPRa normalizes firing activity in *Itgb3* Het cortical networks. **(A)** Top, representative confocal images of primary cortical neurons on MEAs from WT, *Itgb3* Het and KO mice expressing the indicated constructs. Middle, transduction efficiency was confirmed by EGFP expression. Bottom, representative traces from two electrodes. **(B)** Representative raster plots of network activity for experiments as in (A). **(C-G)** Quantification of experiments as in (B). Unlike exogenous expression of β3 integrin, enhancement of endogenous β3 integrin levels by CRISPRa in Het cultures normalizes cortical network activity (*p<0.05, **p<0.01, ***p<0.001, one-way ANOVA followed by Tukey post-test, n=15, 15 and 9 for each WT, Het and KO condition, respectively; 4-5 independent cultures). Data are presented as mean±SEM; dots represent individual values. **(H)** Representative heatmaps of Pearson’s correlation coefficients (*r*) for burst activity from wells containing 9 electrodes (E1-9). Network synchrony correlates with β3 integrin levels **(I)** Quantification of experiment as in (H). All electrode pairs exhibited a positive *r*. The graph shows the percentage of *r* with a p-value <0.05 (Positively correlated) and a p-value >0.05 (Non-correlated). n = 473, 493, 447, 457, 455, 532, 257 and 279 pairs for WT+gRNA Ctrl, WT+gRNAs 2+3, Het+gRNA Ctrl, Het+gRNA 3, Het+gRNAs 2+3, Het+exogenous β3 integrin, KO+gRNA Ctrl and KO+gRNAs 2+3, respectively (*p<0.05, **p<0.01, ***p<0.001, Chi-square test). See also Fig S3.

CRISPRa restored Het but not KO activity to WT values (**Fig 3A-C** and **F**), thus confirming that the effects on network activity were due specifically to β3 integrin. Exogenous expression of this protein enhanced instead firing and burst rate well above WT values (**Fig 3A-G**).

Blockade of GABAergic inhibition with bicuculline increased all parameters of firing activity in all experimental conditions (**Fig S3I-O**). Some effects on neurons expressing exogenous β3 integrin were less pronounced, possibly because of a ceiling effect (**Fig S3L, N**).

Simultaneous occurrence of bursts across a network is indicative of functional interconnected neurons (Bikbaev et al., 2015). Notably, burst synchrony was lower in networks of Het and KO neurons than WT neurons. While CRISPRa could largely reverse burst correlation in Het, exogenous β3 integrin induced over-synchronous bursting (**Fig 3H, I**).

Collectively, these findings indicate that fine-tuning of β3 integrin expression can be used to regulate network activity and synchrony within a physiological range.

### β3 integrin coordinates neuronal activity in primary cortical neurons

To monitor spontaneous network activity at the level of individual neurons, we used the Ca^2+^indicator jRCaMP1b. Somatic fluorescence transients exhibited differences in frequency between genotypes and rescue conditions similar to those found for the firing rate in the MEA recordings (**Fig 4A, B** vs. **Fig 3A-C**).

**Fig 4.**
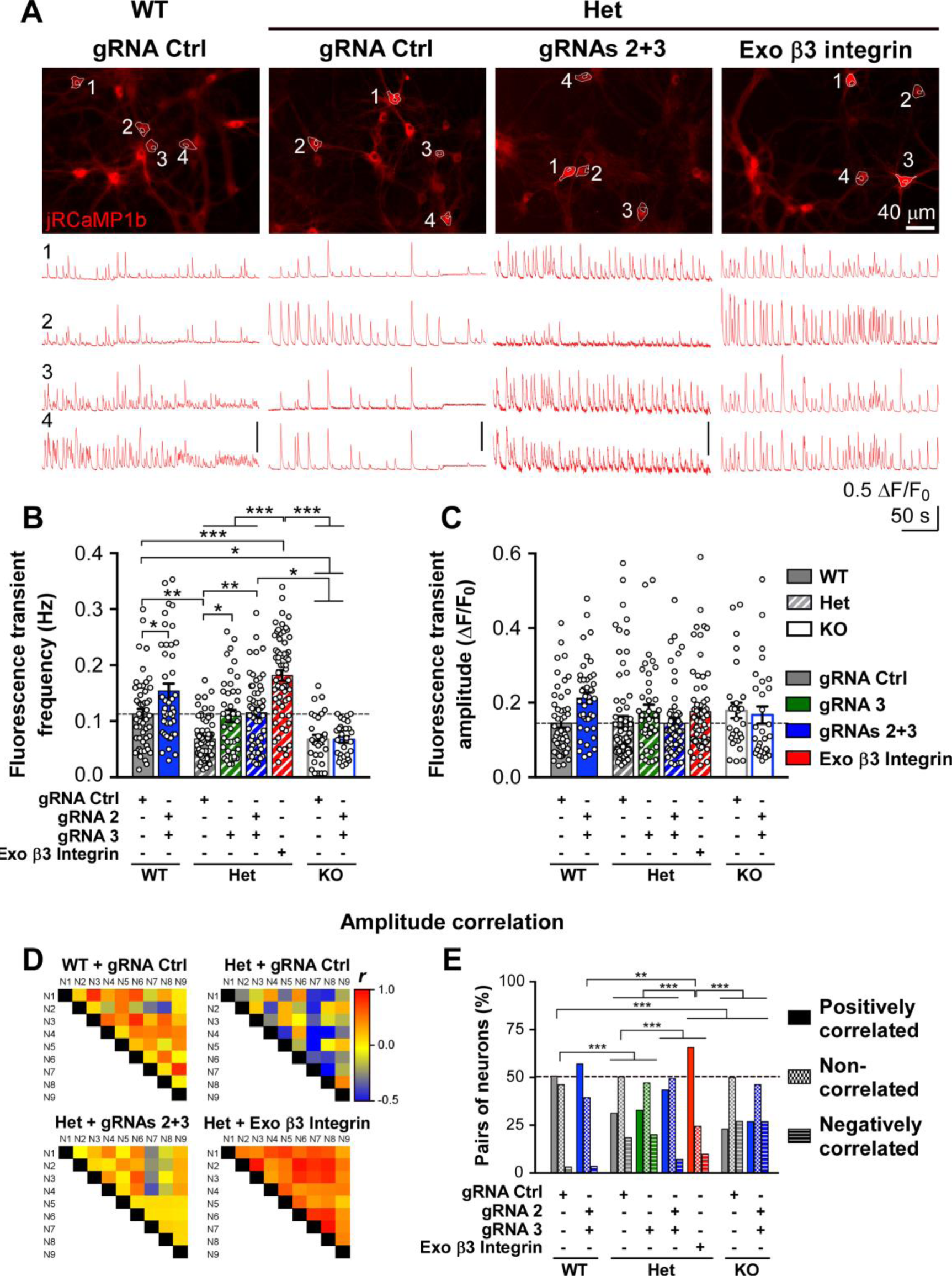
CRISPRa normalizes spontaneous activity in *Itgb3* Het neurons. **(A)** Top, representative images of jRCaMP1b in primary cortical cultures from WT and *Itgb3* Het mice expressing the indicated constructs. Images are an average over 5 min recording period. Bottom, traces show spontaneous somatic fluorescence responses from the indicated ROIs. **(B-C)** Quantification of experiments as in (A) for frequency (B) and amplitude (C). Enhancement of endogenous β3 integrin levels by CRISPRa rescues the reduced frequency of fluorescence transients in Het cultures (*p<0.05, **p<0.01, ***p<0.001, one-way ANOVA followed by Tukey post-test, n=50, 43, 69, 40, 54, 63, 29 and 29 for WT+gRNA Ctrl, WT+gRNAs 2+3, Het+gRNA Ctrl, Het+gRNA 3, Het+gRNAs 2+3, Het+exogenous β3 integrin, KO+gRNA Ctrl and KO+gRNAs 2+3, respectively; 4-5 independent cultures). Data are shown as mean±SEM; dots represent individual values. **(D)** Heatmaps of Pearson’s correlation coefficients (*r*) for fluorescence transient amplitudes of nine neurons (N1-9) from representative fields of view. The *r* for amplitude of fluorescence signals correlates with β3 integrin levels. **(E)** Quantification of experiment as in (D). The graph shows the percentage of positive *r* with a p-value <0.05 (Positively correlated), negative *r* with a p-value <0.05 (Negatively correlated) and *r* with a p-value >0.05 (Non-correlated). n=158, 114, 396, 125, 168, 233, 48 and 119 pairs for WT+gRNA Ctrl, WT+gRNAs 2+3, Het+gRNA Ctrl, Het+gRNA 3, Het+gRNAs 2+3, Het+exogenous β3 integrin, KO+gRNA Ctrl and KO+gRNAs 2+3, respectively (**p<0.01, ***p<0.001, Chi-square test). See also Fig S4.

While MEAs provided information mainly in the temporal domain, jRCaMP1b fluorescence transients exhibited high variability in amplitude also within the same recordings (**Figs 1A** and **4A**), suggesting variability in the underlying firing activity (Inoue et al., 2019). Although the average amplitude of the fluorescence signals was not different across conditions (**Fig 4C**), their variability allowed us to gain a better insight into the level of synchrony of the networks. Because a positive correlation in the amplitude profiles between two neurons is indicative of similar firing activities (**Fig S4A**), we compared the fluorescence amplitude profiles of neurons within a network (**Fig 4D, E**). 51, 31 and 23% of neuron pairs were positively correlated in WT, Het and KO networks, respectively (**Fig 4D, E** and **S4B**). In Het, CRISPRa could restore the percentage of positively correlated neuron pairs to WT values while expression of exogenous β3 integrin increased positive correlation above WT levels.

Altogether, these findings indicate that β3 integrin is critical to coordinating cortical neuronal activity.

## DISCUSSION

We propose that β3 integrin positively regulates network excitability in cortical primary neurons and that it does so by increasing the level of synchronous neuronal firing. We have demonstrated this using two complementary approaches: MEA recordings and Ca^2+^ imaging of populations of neurons in cortical networks from WT, *Itgb3* Het and KO mice. Further, we have designed CRISPRa tools to titrate expression of endogenous β3 integrin, thus overcoming the limitations posed by overexpression of exogenous genes.

A large body of evidence indicates that multiple cues from ECM and glial cells regulate network activity. Enzymatic digestion of ECM components induces epileptiform activity in primary neuronal cultures (Bikbaev et al., 2015; Korotchenko et al., 2014; Vedunova et al., 2013). Integrins on the neuronal surface are ideally positioned to adjust neuronal activity in response to changes in the ECM. For instance, the glia-released factors TNFα and SPARC increase and decrease expression of β3 integrin, respectively (Cingolani et al., 2008; Jones et al., 2011). However, whether these changes affect neuronal function has not been addressed.

We have previously shown that there are no major differences in spontaneous excitatory synaptic transmission between WT, *Itgb3* Het and KO cultures under basal conditions and that an increase in excitatory synaptic transmission becomes apparent only when β3 integrin is overexpressed (Cingolani et al., 2008). Here, we used primary cortical cultures, which generate spontaneous neuronal networks, to address whether changes in β3 integrin expression within a physiological range affect the overall activity of the network. We find that increasing β3 integrin levels with CRISPRa enhances network excitability (**Fig 3** and **4**). Because surface expression of this integrin is negatively regulated by neuronal activity (Cingolani et al., 2008), we propose the existence of a negative feedback loop between β3 integrin and network excitability.

Simultaneous recordings from multiple sites provide the possibility to determine whether the neurons in the network fire independently of each other or in synchrony. Both MEA recordings and Ca^2+^ imaging indicated that β3 integrin expression tightly correlates with network synchrony (**Fig 3H, I, 4D** and **E**). Several factors determine network dynamics, such as the number of shared connections, the excitatory/inhibitory ratio and the level of intrinsic excitability (Hahn et al., 2019; Silberberg et al., 2005). β3 integrin positively modulates excitatory synaptic strength (Cingolani and Goda, 2008; Cingolani et al., 2008; Pozo et al., 2012) but not the overall dendritic density of excitatory synaptic connections (**Fig S3C, G**). To clarify whether changes in inhibitory drive also contribute to the observed differences in network dynamics, we blocked fast inhibitory synaptic transmission (**Fig S3I-O**). As expected, disinhibition boosted firing and burst rate, though to a similar extent in most conditions considered. Hence, the strength of excitatory synaptic coupling seems to play a major role in the β3 integrin-dependent effects on network excitability and synchrony. Although the present findings have been obtained in a simplified culture system devoid of sensory inputs, it is worth noting that reduced neural synchronization may underlie some aspects of ASD defects in network signaling propagation (Uhlhaas et al., 2009).

A significant increase in copy number variations has been reported in ASD patients (Glessner et al., 2009; Levy et al., 2011; Pinto et al., 2010; Sebat et al., 2007). Further, the majority of ASD mutations affect a single allele (de la Torre-Ubieta et al., 2016). These observations illustrate the importance of gene dosage in ASD. It would therefore be highly desirable to find effective strategies for rebalancing gene deficiency in ASD and, more in general, in brain disorders.

Here, we designed CRISPRa tools to activate *Itgb3*, which is deficient in some cases of ASD (Cantor et al., 2005; Carter et al., 2011; Dohn et al., 2017; O’Roak et al., 2012; Weiss et al., 2006), and demonstrated that *Itgb3* haplo-insufficiency can be precisely compensated in neurons by activating the transcription of the remaining functional allele. By contrast, even if we used mild over-expression conditions, as given by a lentivirus with the relatively weak and developmentally-delayed Synapsin promoter, these led to a ∼50-fold increase in β3 integrin protein levels, aberrant protein localization, and hyperactive and over-synchronous cortical networks (**Figs 2, 3,** and **4**). Thus, activation of endogenous genes by CRISPRa reflects a more natural mechanism of action and, whenever possible, should be preferred to traditional overexpression of exogenous genes in rescue experiments. Although CRISPRa holds great promise for diseases caused by haplo-insufficiency, it is nevertheless not suitable for compensating dominant mutations as it will increase expression of both functional and aberrant alleles.

Different degrees of dCas9-dependent gene activation can be achieved using different transcriptional activators (Chavez et al., 2015; Konermann et al., 2015; Perez-Pinera et al., 2013; Tanenbaum et al., 2014; Zhou et al., 2018). We chose VP64 because it has a small size and a moderate activation potential (Chavez et al., 2015; Matharu et al., 2019). This allowed us to use a single lentiviral vector to drive expression of both dCas9-VP64 and a gRNA in neurons and to obtain physiologically relevant β3 integrin dosage levels.

The CRISPRa system is not as prone as CRISPR/Cas9 to off-target effects because it does not cleave chromosomal DNA. Spurious activation of off-target genes would require dCas9-VP64 to bind specifically to promoter regions. Indeed, previous studies have failed to identify significant off-targets for CRISPRa in neurons (Matharu et al., 2019; Savell et al., 2019). Similarly, using ChIP-qPCR, we identified weak binding of dCas9 to only 10% of the predicted off-target sites without however any detectable effect on gene expression (**Fig S2**). The absence of physiological effects of CRISPRa in *Itgb3* KO neurons further confirmed the specificity of the system (**Fig 3** and **4**). Our results together with the findings from previous studies suggest therefore that CRISPRa has high specificity in neurons.

In summary, β3 integrin positively regulates cortical network activity and synchrony. Further, titration of this protein by CRISPRa provides a precise and efficient system for modulating neuronal network activity.

## ACKNOWLEDGMENTS

We thank F Benfenati (IIT) for support, D. Moruzzo and E. Colombo (IIT) for technical help. This work was supported by the Compagnia San Paolo (Proposal ID: 2015 0702 to L.A.C.) and by IIT.

## AUTHOR CONTRIBUTIONS

F.J., A.T. and L.C. designed and performed experiments, analyzed data and wrote the manuscript.

## DECLARATION OF INTERESTS

The authors declare no competing interests.

## METHODS

### CONTACT FOR REAGENT AND RESOURCE SHARING

Further information and requests for resources and reagents should be directed to and will be fulfilled by the Lead Contact, Lorenzo Cingolani.

### EXPERIMENTAL MODEL AND SUBJECT DETAILS

All experiments were performed in accordance with EU and Italian regulation. Itgb3 KO mice (B6;129S2-Itgb3tm1Hyn /J, Jackson Laboratory) were described previously (Cingolani et al., 2008; McGeachie et al., 2012) and were backcrossed to the C57BL/6 background >10 times at the time of experiments.

## METHOD DETAILS

### gRNA design and plasmid construction

We used the web tool http://crispr.mit.edu/ (Ran et al., 2013), which maximizes the regions with low off-target probability, to design three gRNAs targeting the region from -100 to -200 bp relative to the transcription start site (TSS) of the *Itgb3* gene. As negative control (gRNA Ctrl), we used a non-targeting gRNA sequence (**Fig S1F**). The pU6-(BbsI)-EF1a-dCas9-VP64-T2A-EGFP plasmid (**Fig S1F**), used to co-express a gRNA, the dCas9-VP64 fusion protein and EGFP, was constructed by inserting the EF1a-dCas9-VP64-T2A-EGFP cassette from the dCAS9-VP64-GFP plasmid (gift from Feng Zhang; Cat. No. 61422, Addgene) (Konermann et al., 2015) in place of the CBh-Cas9-T2A-mCherry cassette of the pU6-(BbsI)-CBh-Cas9-T2A-mCherry plasmid (gift from Ralf Kuehn; Cat. No. 64324, Addgene) (Chu et al., 2015). The gRNA sequences were inserted downstream of the U6 promoter using the BbsI cloning sites.

The lentiviral vectors pLL-U6-(gRNA)-EF1a-dCas9-VP64-T2A-EGFP (**Fig 2A**) were constructed by inserting the cassettes U6-gRNA Ctrl, U6-gRNA 2 or U6-gRNA 3 from the pU6-(gRNA)-EF1a-dCas9-VP64-T2A-EGFP plasmids described above in dCAS9-VP64-GFP (Cat. No. 61422, Addgene) using the PacI and AgeI sites. Human β3 integrin was expressed under the control of the short human Synapsin promoter using the lentiviral vector pLL-Syn-EGFP-P2A-ITGB3 (McGeachie et al., 2012). Constructs were generated by standard cloning strategies and verified by sequencing.

### N2a cell culture and transfection

N2a mouse neuroblastoma cells were cultured in Dulbecco’s Modified Eagle Medium (DMEM, Gibco) supplemented with 10% FBS, 2 mM glutamine, 100 U/ml penicillin and 0.1 mg/ml streptomycin (complete culture medium), and maintained in a 5% CO_2_ humidified incubator at 37°C.

Transfection was performed in 60-70% confluent cultures seeded in 6-well plates at 200,000 cells/well in complete culture medium the previous day. Cells were transfected with 3 μg DNA/well using the Ca^2+^ phosphate method (Thalhammer et al., 2018) and used 24-48 hours post-transfection.

### Cell adhesion assay

N2a cells were trypsinized two days after transfection and seeded in complete culture medium on fibronectin-coated coverslips (5 μg/ml for 16 hours; Cat. No. F8141, Sigma) in 24-well plates at a density of 100,000 cells/well. After 1 hour at 37°C, coverslips were washed 4 times with PBS to remove non-attached cells; remaining adherent cells were fixed in 4% PFA, stained with Hoechst and mounted with ProLong Gold mounting medium (ThermoFisher Scientific). To quantify the number of attached cells, three images per each coverslip were taken using a Leica SP8 confocal microscope with a 40x oil immersion objective (NA 1.30); for each condition, six coverslips from three independent cultures were imaged in total.

### Primary cortical culture

Cortical neuronal cultures were prepared from P0 *Itgb3*^*+/+*^(WT), *Itgb3*^*+/*−^(Het) or *Itgb3*^−*/*−^(KO) pups as previously described (Thalhammer et al., 2017), with minor modifications. Briefly, cortices were dissected in ice-cold HBSS, digested with papain (30 U; Cat. No. 3126, Worthington) for 40 min at 37°C, washed and triturated in attachment medium (BME medium supplemented with 10% FBS, 3 mg/ml glucose, 1 mM sodium pyruvate and 10 mM HEPES-NaOH [pH 7.40]) with a flame-polished glass Pasteur pipette. For RT-qPCR and Western blot experiments, cells were seeded at a concentration of 750,000 cells/well onto 6-well plates coated with 2.5 μg/ml poly-D-lysine (PDL; P7405, Sigma) and 1 μg/ml laminin (L2020, Sigma); for confocal microscopy and Ca^2+^imaging experiments, cells were seeded at 75,000/well onto 1.2 cm diameter glass coverslips coated with PDL/laminin as above. After 4 hours, the attachment medium was replaced with maintenance medium (Neurobasal medium supplemented with 2.6% B27, 6 mg/ml glucose, 2 mM GlutaMax, 90 U/ml penicillin and 0.09 mg/ml streptomycin). To prevent glia overgrowth, 0.5 μM of cytosine β-D-arabinofuranoside (AraC) was added at 5 DIV.

### Lentivirus production and infection

HEK293T cells were maintained in Iscove’s Modified Dulbecco’s Medium supplemented with 10% FBS, 2 mM Glutamine, 100 U/ml penicillin and 0.1 mg/ml streptomycin in a 5% CO_2_ humidified incubator at 37°C. Cells were transfected with the Δ8.9 encapsidation plasmid, the VSVG envelope plasmid and the pLL-U6-(gRNA)-EF1a-dCas9-VP64-T2A-EGFP or the pLL-Syn-EGFP-P2A-ITGB3 plasmids described above using the Ca^2+^ phosphate method. The transfection medium was replaced by fresh medium after 14 hours. Supernatants were collected 36 to 48 hours after transfection, centrifuged to remove cell debris, passed through a 0.45 μm filter and ultra-centrifuged two hours at 20,000 g at 4°C. Viral pellets were re-suspended in PBS, aliquoted and stored at -80°C until use (Thalhammer et al., 2018). Neuronal cultures were infected at 6 DIV with the lowest infectious dose capable of transducing ≥95% of neurons (dilution range: 1:300 to 1:700) and used for experiments after ≥10 days (**Fig 2A**).

### Western blotting

Membrane protein-enriched fractions were prepared from cortical neurons at 16 DIV as previously described (Thalhammer et al., 2018). Briefly, cells were washed once in ice-cold PBS and scraped in 100 μl buffer A (25 mM Tris-HCl pH 7.4, 150 mM NaCl, 2 mM KCl, 2.5 mM EDTA) supplemented with protease and phosphatase inhibitors (complete EDTA-free protease inhibitors [Cat. No. 1187358001, Roche Diagnostic]; serine/threonine and tyrosine phosphatase inhibitor, [Cat. No. P0044 and P5726, Sigma]). After removal of the cell debris at 1,000 x g, 4°C for 10 min, the supernatant was centrifuged at 15,000 x g, 4°C for 15 min. The resulting pellet was dissolved in 100 μl RIPA buffer (50 mM Tris pH 8.0, 150 mM NaCl, 1% NP-40, 0.5% sodium deoxycholate, 0.1% SDS) and centrifuged at 15,000 x g, 4°C for 15 min. The resulting supernatant was used for Western blot analysis. Protein concentration was quantified with the BCA Protein Assay kit (Cat. No. 23227, ThermoFisher Scientific). Proteins were separated by SDS-PAGE using 7.5% acrylamide gels and transferred on PVDF membranes. After incubation with primary rabbit anti-integrin β3 (1:200; Cat. No. 4702, Cell signaling) or rabbit anti-β-tubulin III (1:1000; Cat. No. T2200, Sigma) antibodies, membranes were incubated with secondary HRP-conjugated goat anti-rabbit antibody (1: 5000; Cat. No. 31460, ThermoFisher scientific) and immunocomplexes were detected with the chemiluminescent substrate (Cat. No. RPN2106, ECL Prime Western Blotting System, GE Healthcare). We acquired chemiluminescent signals using a ChemiDoc imaging system (Biorad) and quantified immunoreactive bands using ImageJ (http://rsb.info.nih.gov/ij). Band intensity from different samples was normalized to that of WT control within the same membrane.

### RNA extraction and RT-qPCR

Total RNA was extracted with QIAzol lysis reagent (Cat. No. 79306, Qiagen) from primary cortical cultures at 16 DIV as previously described (Thalhammer et al., 2018). We prepared cDNAs by reverse transcription of 1 μg of RNA using the QuantiTect Reverse Transcription Kit (Cat. No. 205311, Qiagen). RT-qPCR was performed in triplicate with 10 ng of template cDNA using iQ™ SYBR^®^ Green Supermix (Cat. No. 1708886, Biorad) on a CFX96 Real-Time PCR Detection System (Biorad) with the following universal conditions: 5 min at 95 °C, 45 cycles of denaturation at 95 °C for 15 sec, and annealing/extension at 60 °C for 45 sec. Primers were designed with Primer-BLAST (https://www.ncbi.nlm.nih.gov/tools/primer-blast) to avoid significant cross homology regions with other genes. Product specificity and absence of primer dimers was verified by melting curve analysis and agarose gel electrophoresis. qPCR reaction efficiency for each primer pair was calculated by the standard curve method with a five points serial dilution of cDNA. Calculated qPCR efficiency for each primer set was used for subsequent analysis. The relative quantification of gene expression was determined using the ΔΔCt method. Data were normalized to glyceraldehyde-3-phosphate dehydrogenase (GAPDH), β-actin (ACTB) and hypoxanthine phosphoribosyltransferase 1 (HPRT1) by the multiple internal control gene method with GeNorm algorithm (Vandesompele et al., 2002). mRNA expression was normalized to WT control samples within the same RT-qPCR plate. Sequences of all the primers used are listed in the STAR Methods Key Resources Table.

### Chromatin immunoprecipitation

After cross-linking with 1% formaldehyde for 10 min and quenching with 125 mM glycine for 5 min, we extracted the chromatin from transduced primary cortical neurons using the Chromatin Extraction kit (Cat. No. ab117152, Abcam) according to the manufacturer’s instructions. After sonication, samples were further processed for chromatin immunoprecipitation (IP) using the ChIP Kit-One step (Cat. No.ab117138, Abcam) and a rabbit polyclonal anti-Cas9 antibody (Cat. No. C15310258, Diagenode) or a control non-immune IgG (Cat. No. ab117138, Abcam; **Fig S2B**). Enrichment of target regions was assessed by RT-qPCR as detailed in previous section using the primers listed in the STAR Methods Key Resources Table.

### Confocal microscopy and image analysis

Ten days post-infection, cultures were fixed for 8 min with 4% PFA/4% sucrose at room temperature (RT), treated for 10 min at 50°C with a sodium citrate solution (10 mM tri-sodium citrate dihydrate, pH 6.0, 0.05% Tween-20) to retrieve the β3 integrin antigen (Mazalouskas et al., 2015) and permeabilized for 10 min at RT with 0.1% TritonX-100. β3 integrin staining was revealed using a rabbit monoclonal anti-integrin β3 (1:200; Cat. No. 13166, Cell signaling) and the Tyramide SuperBoost kit (Cat. No. B40922, ThermoFisher scientific) with the Alexa Fluor568 Tyramide Reagent (10 min, 1:10 dilution; Cat. No. B40956, ThermoFisher scientific) before counterstaining for GFP, vGlut1 and total GluA2 using the following primary antibodies: chicken anti-GFP (1:1000; Cat. No. AB13970, Abcam,), guinea pig anti-vGlut1 (1:500; Cat. No. 135304, Synaptic System) and mouse anti-GluA2 (1:300; Cat. No. MAB397, Millipore), respectively. Secondary antibodies were Alexa Fluor488-conjugated anti-chicken (1:1000; Cat. No. A11039, ThermoFisher scientific), Dylight405-conjugated donkey anti-guinea pig (1:150; Cat. No. 706-475-148, Jackson ImmunoResearch) and Alexa Fluor647-conjugated goat anti-mouse (1:800; Cat. No. A21236, ThermoFisher scientific). Confocal stacks were acquired at 200 Hz with a Leica SP8 using a 63x oil immersion objective (NA 1.40), 1.2x digital zoom, 0.15 μm pixel size, 1 AU pinhole, 0.3 μm between optical sections, with a sequential line-scan mode and 3x scan averaging. For all experimental conditions compared, the same settings for laser intensity, offset and PMT gain were used.

Confocal images were analyzed using ImageJ. Each stack was filtered using a Gaussian filter (radius: 0.5 pixels), and the maximal fluorescence intensities of in-focus stacks were Z-projected. The resulting images were automatically thresholded using the Robust Automatic Threshold Selection plugin followed by the watershed algorithm. Dendritic analysis was performed on dendritic ROIs of 40-120 μm in lengths, manually selected in the GFP channel blind to the experimental condition. Co-localization was estimated for the thresholded ROIs with the Coloc2 plugin using the Manders’ coefficients (M_A_=∑_i_ A_i, coloc_/∑ _i_ A_i,_ where ∑ _i_A_i_ is the sum of intensities of all pixels above threshold for channel A and ∑_i_ A_i,coloc_ is calculated as ∑_i_ A_i_ but only for pixels where also the second channel B is above threshold).

### Multielectrode array recordings

Cortical neurons were seeded at 150,000 cells/well on 6-well multi-electrode arrays (MEAs, Multichannel Systems) coated with PDL/laminin. Each well contained nine electrodes (30 μm diameter; 200 μm center-to-center spacing). Neurons were transduced at 6 DIV and network activity was recorded at 16-17 DIV using a MEA1060INV amplifier (Multichannel Systems). Neurons were kept in maintenance medium at 37°C throughout the recordings. To ensure stabilization of the electrical signal, experiments were initiated 10 min after transferring the MEAs from the incubator to the set-up. Network activity was recorded first under basal conditions (5 min) and subsequently in response to bicuculline application (10 μM; 10 min; **Fig S3I**).

Spike detection and spike train analysis were performed with the MC-Rack software (Multichannel Systems). Spike threshold was set for each electrode at 5 times the standard deviation of the baseline noise level. Bursts were operationally defined as a collection of a minimum number of spikes (N_min_ = 5) separated by a maximum interspike interval (ISI_max_ = 100 ms) (Chiappalone M, 2005). Following the burst detection procedure, several measures describing spike and burst statistics were extracted including mean firing rate, burst rate, mean burst duration, percentage of spike in burst and intraburst spike frequency. The Pearson’s correlation coefficient for burst activity was computed for all electrode pairs in each MEA well.

### Ca^2+^ imaging

Imaging was performed in primary cortical cultures at 30 ± 2°C in aCSF containing (in mM): 140 NaCl, 3.5 KCl, 2.2 CaCl_2_, 1.5 MgCl_2_, 10 D-glucose, and 10 HEPES-NaOH (pH 7.38; osmolarity adjusted to 290 mOsm). In initial experiments, we tested the red-shifted genetically encoded Ca^2+^ indicators (GECIs) jRGECO1a, jRCaMP1a and jRCaMP1b (Dana et al., 2016) and chose jRCaMP1b for subsequent experiments as it provided overall the best signal-to-noise ratio, the largest dynamic range and best temporal resolution of the three GECIs in our experimental conditions. Cultures were infected with the appropriate lentivirus and pAAV.Syn.NES-jRCaMP1b.WPRE.SV40 (Cat. No. 100851-AAV1, Addgene; titer: 1.7*10^13^ GC/ml; dilution 1: 50,000) 9-12 and 4-6 days prior to experiments, respectively, and recorded at 15-18 DIV. Imaging was performed with a cooled charge-coupled device (CCD) camera (ORCA-R2, Hamamatsu) mounted on an inverted microscope (DMI6000B, Leica) with a 20x, 0.75 NA glycerol immersion objective. A 200W metal halide lamp (Lumen200Pro, Prior Scientific) and a filter set comprising a BP 515-560 nm excitation filter, a 580 nm dichroic mirror and a LP 590 emission filter (Filter set N2.1, Leica) were used for illumination. Images were captured at 15.3 Hz with 50 ms integration times at a depth of 8 bits. Network activity was recorded for 5 min.

Images were analyzed in ImageJ with the plugin Time Series Analyzer V3.0 (http://rsb.info.nih.gov/ij/plugins/time-series.html) and with customized routines in Igor Pro 6.03 (www.wavemetrics.com). Regions of interest (ROIs) were manually drawn on the soma (excluding the nuclear region; **Fig 4A**) of each neuron exhibiting at least one spontaneous fluorescence transient above two SD of the background noise during 5 min recording period. The intensity of twin ROIs positioned within 50 μm were used to subtract the background noise. Signals were quantified as ΔF/F_0_, where ΔF=F-F_0_, with F_0_ measured over 1 s period preceding the fluorescence transient. The Pearson’s correlation coefficient for fluorescence transient amplitude was computed for all neuron pairs in each field of view.

### Statistical analysis

Unless otherwise stated, statistical differences were assessed using unpaired two-tailed Student’s t-test and the one-way analysis of variance (ANOVA) test followed by the Tukey-Kramer post-test, as required. The Chi-squared test was used for figures 3I and 4E (Prism 7, GraphPad Software Inc.). Average data are expressed as mean±SEM.

