## Supplemental Figures 1 - 4 for "Correction of β3 integrin haplo-insufficiency by CRISPRa normalizes cortical network activity"

### SUPPLEMENTARY FIGURES

**Figure S1.** **β3 integrin expression in mouse primary cortical neurons and N2a cells.** Related to Figures 1 and 2.

**Figure S2.** **Targeting specificity of CRISPR/dCas9 for β3 integrin in primary cortical neurons.** Related to Figures 2, 3 and 4.

**Figure S3.** **Effects of β3 integrin on synaptic marker expression and network activity under blockade of $GABA_A$-mediated inhibitory transmission.** Related to Figures 2 and 3.

**Figure S4.** **Further characterization of the amplitude correlation of fluorescence transients and graphical summary.** Related to Figures 1-4.

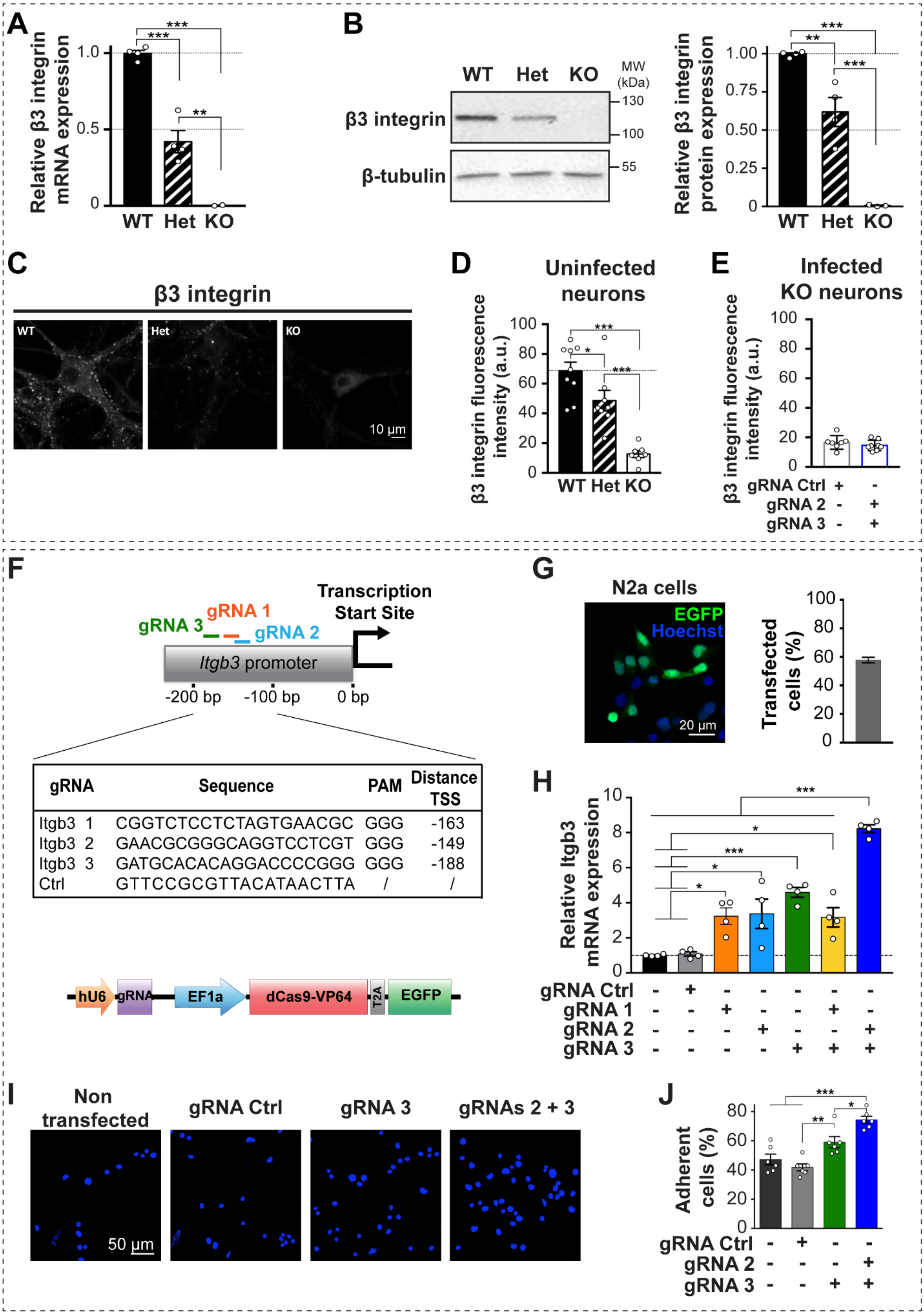
A
Relative β3 integrin mRNA expression
1.0
0.5
0
***
**
WT Het KO
B
WT Het KO
MW (kDa)
β3 integrin
β-tubulin
130
100
55
Relative β3 integrin protein expression
1.00
0.75
0.50
0.25
0
C
β3 integrin
WT
Het
KO
10 µm
D
Uninfected neurons
β3 integrin fluorescence intensity (a.u.)
100
80
60
40
20
0
*
***
WT Het KO
E
Infected KO neurons
gRNA Ctrl + -
gRNA 2 - +
gRNA 3 - +
F
Transcription Start Site
gRNA 1
gRNA 2
gRNA 3
Itgb3 promoter
-200 bp
-100 bp
0 bp
gRNA | Sequence | PAM | Distance TSS
Itgb3 1 | CGGTCTCCTCTAGTGAACGC | GGG | -163
Itgb3 2 | GAACGCGGGCAGGTCCTCGT | GGG | -149
Itgb3 3 | GATGCACACAGGACCCCGGG | GGG | -188
Ctrl | GTTCCGCGTTACATAACTTA | / | /
hU6
gRNA
EF1a
dCas9-VP64
T2A
EGFP
G
N2a cells
EGFP
Hoechst
20 µm
Transfected cells (%)
100
80
60
40
20
0
H
Relative Itgb3 mRNA expression
10
8
6
4
2
0
***
*
gRNA Ctrl - + - - - - -
gRNA 1 - - + - - + -
gRNA 2 - - - + - - +
gRNA 3 - - - - + + +
I
Non transfected
gRNA Ctrl
gRNA 3
gRNAs 2 + 3
50 µm
J
Adherent cells (%)
80
60
40
20
0
***
**
*
gRNA Ctrl - + - -
gRNA 2 - - - +
gRNA 3 - - + +

**Fig S1. β3 integrin expression in mouse primary cortical neurons and N2a cells. Related to Figs 1 and 2. (A)** Expression of β3 integrin mRNA in mouse primary cortical neurons at 16 DIV. Values are normalized to WT samples within the same RT-qPCR plates (n=4, 4 and 2 independent cultures for WT, *Itgb3* Het and KO, respectively). **(B)** Membrane protein fractions from mouse primary cortical neurons were analysed by Western blotting at 16 DIV. Left, representative immunoblots for β3 integrin, β-tubulin was used as loading control; right, quantification of immune-reactive bands; band intensities were normalized to WT within the same membrane (n=4, 4 and 3 independent cultures for WT, Het and KO, respectively). **(C)** Representative confocal images of WT, *Itgb3* Het and KO primary cortical neurons stained for β3 integrin at 16 DIV. **(D)** Quantification of experiments as in (C; n=9, 8 and 7 images for WT, Het and KO, respectively). **(E)** Quantification of experiments as in Figs 2E, F. Infection with CRISPRa and gRNAs 2+3 does not increase β3 integrin expression in *Itgb3* KO cultures (n=7 and 8 images for gRNA Control and gRNAs 2+3, respectively). **(F)** Top, gRNA sequences and position of their targets on the *Itgb3* promoter. Bottom, construct used for transfecting murine N2a cells, containing a cassette for expressing gRNA and one for expressing dCas9-VP64 and EGFP. **(G)** Representative image of transfected N2a cells (left) and quantification of transfection efficiency (right). **(H)** Quantification of β3 integrin mRNA levels in N2a cells 24 hours after transfection with the indicated constructs. mRNA expression was normalized to the values of non-transfected samples within the same RT-qPCR plate (n=4 independent cultures each; 2 technical replicates per culture). **(I-J)** Cell adhesion assay for N2a cells transfected with the indicated constructs and plated onto fibronectin-coated coverslips. Representative images of the adherent cells stained with Hoechst (I) and quantification of the percentage of adherent cells (J; n=6 each from 3 independent cultures). Data are presented as mean±SEM; dots represent individual values (*p<0.05, **p<0.01, ***p<0.001, one-way ANOVA followed by Tukey post-test for panels A, B, D, H and J; p=0.37, unpaired Student's t-test for panel E).

**A**

| | Off-target sequences | Number of mismatches | Genomic coordinates (GRCm38.p4 C57BL/6J) | Position | Gene |
|---|---|---|---|---|---|
| **gRNA 2** | **GAACGCGGGCAGGTCCTCGT** | | | | |
| #1 | GAGCGCTGGCAGGTCCTCGG | 3 | Chr5: 128607080 - 128607099 | Intergenic | |
| #2 | GGATGGGGGAAGGTCCTCGT | 4 | Chr14: 65726265 - 65726284 | Intronic | Scara5 |
| #3 | GACCGCAGAGAGGTCCTCGT | 4 | Chr7: 135803303 - 135803322 | Intergenic | |
| #4 | GATGGCAGGCAGTTCCTCGT | 4 | Chr6: 86380681 - 86380700 | Intergenic | |
| #5 | GGAGGGGGGCAGTTCCTCGT | 4 | Chr4: 44576073 - 44576092 | Intronic | Pax5 |
| #6 | GAACTCCGACCGGTCCTCGT | 4 | Chr13: 34002574 - 34002593 | Exonic | Serpinb6a |
| #7 | AAACCCGGGAAGGTCCTAGT | 4 | Chr16: 89832908 - 89832927 | Intronic | Tiam1 |
| #8 | GAAAGAGTGCAGGTCCTAGT | 4 | Chr11: 60178341 - 60178360 | Intergenic | |
| #9 | GAACACAGGCAGTTCCTCGG | 4 | Chr14: 64286180 - 64286199 | Intronic | Msra |
| #10 | GACCGAGGGCTGGTCCTCAT | 4 | Chr2: 58176765 - 58176784 | Exonic | Gm13546 |
| **gRNA 3** | **GATGCACACAGGACCCCGGG** | | | | |
| #1 | GATGCCCACAGGGCCCCGGG | 2 | Chr4: 108941267 - 108941286 | Exonic | Rab3B |
| #2 | CCTGAACGCAGGACCCCGGG | 4 | Chr12: 105964060 - 105964079 | Intergenic | |
| #3 | CTTGCACACAGGACCCTGGG | 3 | Chr6: 146208088 - 146208107 | Intronic | Itpr2 |
| #4 | GGAGAAGACAGGACCCCGGG | 4 | Chr17: 68274174 - 68274193 | Intergenic | |
| #5 | CCTGTACACAGCACCCCGGG | 4 | Chr8: 122480377 - 122480396 | Exonic | Ctu2 |
| #6 | GCTGAACTCAGCACCCCGGG | 4 | Chr6: 39621848 - 39621867 | Intronic | Braf |
| #7 | GATGAACAGAGGACCCGGGG | 3 | Chr17: 45669894 - 45669913 | Exonic | Tmem63B |
| #8 | GACACACCCAGGACCCCGGA | 4 | Chr1: 91602261 - 91602280 | Intergenic | |
| #9 | GAGGTACTCAGGACCCTGGG | 4 | Chr3: 134181351 - 134181370 | Intergenic | |
| #10 | CCTACACACAGGCCCCCGGG | 4 | Chr12: 117153148 - 117153167 | Intronic | Ptprn2 |

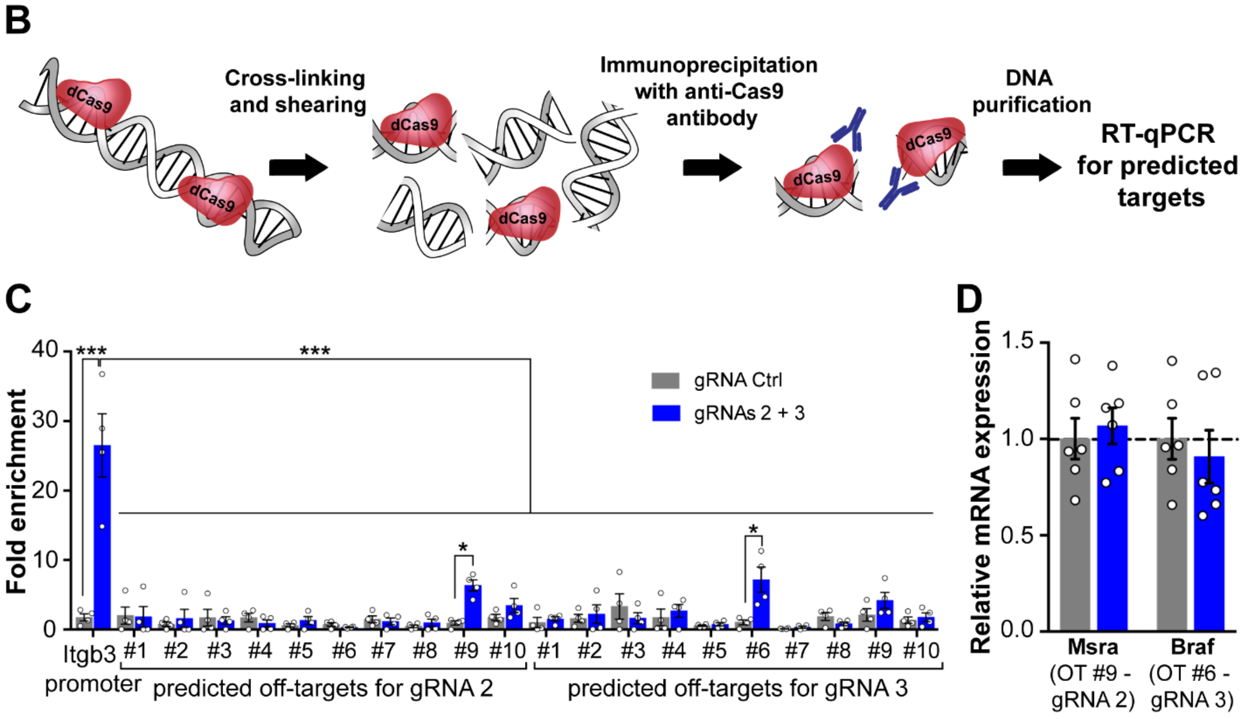


**Fig S2. Targeting specificity of CRISPR/dCas9 for β3 integrin in primary cortical neurons. Related to Figs 2, 3 and 4 (A)** List of top-ten predicted off-targets for gRNA 2 and gRNA 3 (http://crispr.mit.edu and https://crispr.cos.uni-heidelberg.de). The mismatches between the predicted off-targets and the on-target sequence are highlighted in red. Number of mismatches, genomic coordinates, position and name of potentially targeted genes are indicated. Scara5, scavenger receptor class A, member 5; Pax5, paired box protein 5; Serpinb6a, serine/cysteine peptidase inhibitor, clade B, member 6a; Tiam1, T cell lymphoma invasion and metastasis 1; Msra, mitochondrial peptide methionine sulfoxide reductase;

Gm13546, predicted gene 13546, long non-coding RNA; Rab3B, member RAS oncogene family Rab3b; Itpr2, inositol 1,4,5-triphosphate receptor 2, transcript variant 2; Ctu2, cytosolic thiouridylase subunit 2; Braf, Braf transforming gene; Tmem63B, transmembrane protein 63b; Ptprn2, protein tyrosine phosphatase, receptor type, N polypeptide 2. **(B)** ChIP-qPCR workflow. dCas9 co-expressed with gRNA Ctrl or gRNAs 2+3 is allowed to bind to chromatin. After cross-linking, the chromatin is sheared, immune-precipitated with an anti-Cas9 antibody and subjected to RT-qPCR. **(C)** Fold enrichment of dCas9 at on- and predicted off-target sites was calculated over an IgG control IP (*$p<0.05$, ***$p<0.001$, two-way ANOVA followed by Tukey post-test, n=4 independent cultures, 2 technical replicates each). **(D)** RT-qPCR quantification of mRNA expression for the two predicted off-target genes displaying significant dCas9 binding. Expression of both genes is not altered by dCas9-VP64 binding ($p \geq 0.60$, unpaired student's t-test, n=6 independent cultures, 2 technical replicates each). Data are presented as mean±SEM; dots represent individual values.

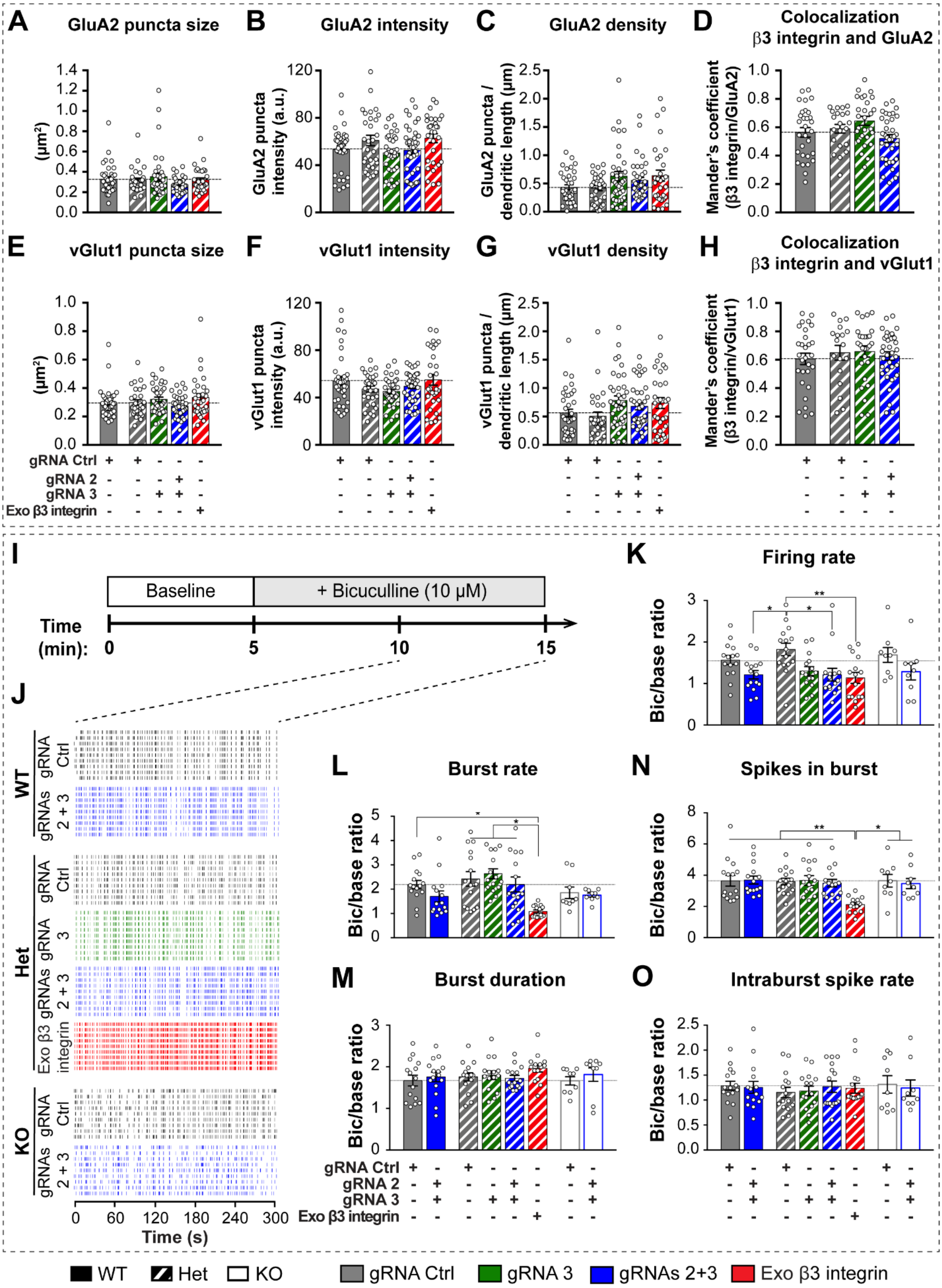


**Fig S3. Effects of β3 integrin on synaptic marker expression, and network activity upon blockade of GABA$_A$-mediated inhibitory transmission. Related to Figs 2 and 3. (A-D)** Quantification of the effects of the indicated gRNAs on dendritic puncta size (A), intensity (B)

and number (C) for total GluA2 and on Mander's co-localization coefficient of β3 integrin with GluA2 (D) in experiments as in Fig 2E. **(E-H)** as in (A-D) but for vGlut1 (n=13, 10, 11, 13 and 10 from 3 independent cultures each for WT+gRNA Ctrl, Het+gRNA Ctrl, Het+gRNA 3, Het+gRNAs 2+3 and Het+exogenous β3 integrin, respectively). **(I)** Experimental timeline for bicuculline application in MEA experiments. **(J)** Representative raster plots of network activity from WT, *Itgb3* Het and KO cultures expressing the indicated constructs. **(K-O)** Quantification of experiments as in (J) for firing rate (K), burst rate (L), burst duration (M), percentage of spikes in burst (N) and intra-burst spike rate (O) expressed relative to the corresponding baseline values (reported in Fig 3) for each recording (* $p<0.05$, ** $p<0.01$, one-way ANOVA followed by Tukey post-test, n=15, 15 and 9 for each WT, Het and KO condition, respectively; 5 independent cultures). Data are presented as mean±SEM; dots represent individual values.

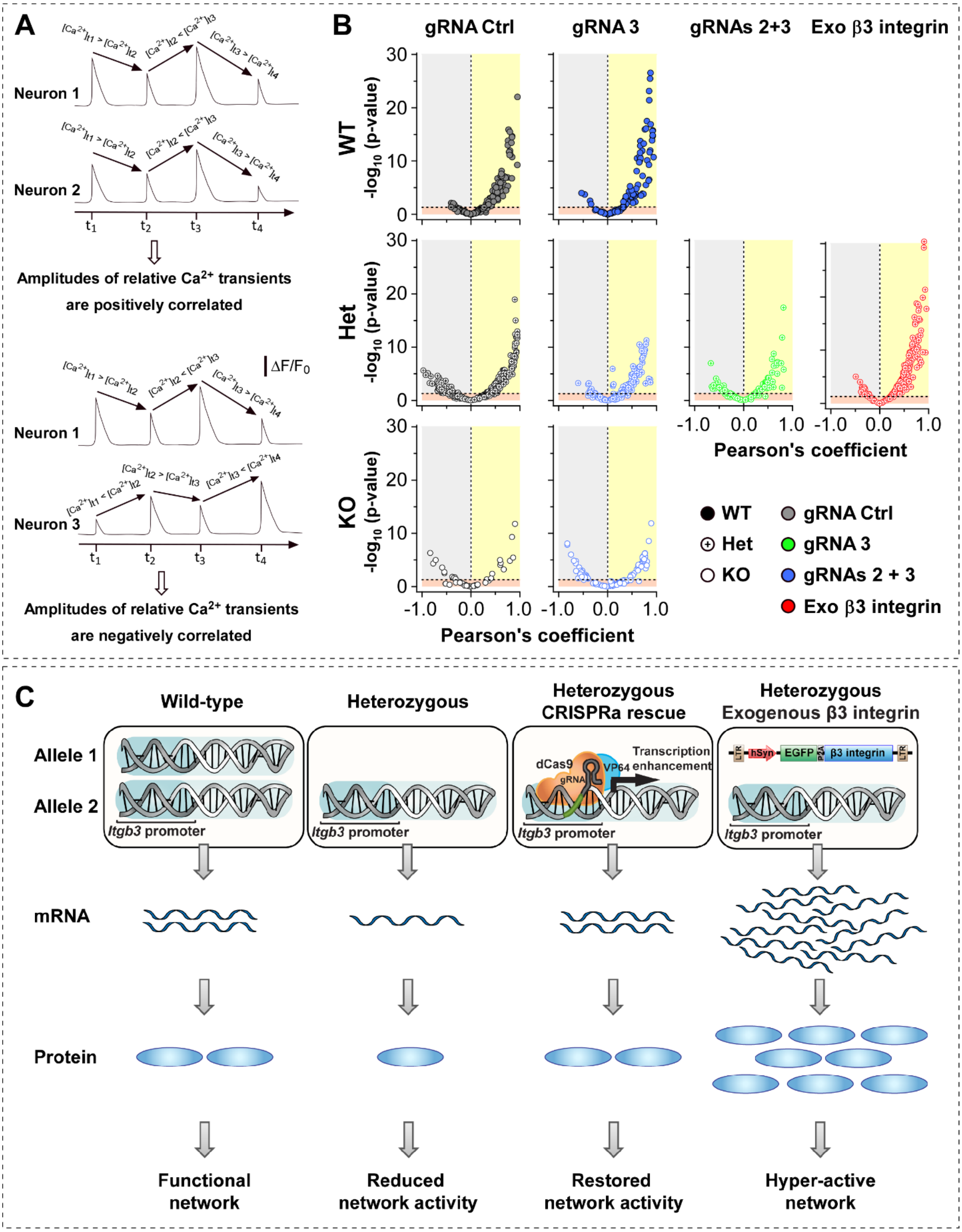


**Fig S4. Further characterization of the amplitude correlation of fluorescence transients and graphical summary. Related to Figs 1-4. (A)** Assuming steady state conditions during the 5-min-long recording period, differences in amplitude of florescence signals are indicative of relative differences in $Ca^{2+}$ transients at different time points within one neuron. Although it is not possible to compare directly differences in amplitude of fluorescence transients across neurons because of differences in jRCaMP1b expression, fluorescence amplitude profiles between pairs of neurons can be compared to reveal positive (top) or negative (bottom) correlation in $Ca^{2+}$ transients. **(B)** Volcano plots of all data points used for Fig 4E. The Pearson's correlation coefficient of fluorescence transient amplitudes for pairs of neurons is plotted against the $-\log_{10}$ of its p-value. Grey, pink and yellow backgrounds indicate negative, non-significant and positive correlation, respectively (n=158, 114, 396, 125, 168, 233, 48 and 119 pairs for WT+gRNA Ctrl, WT+gRNAs 2+3, Het+gRNA Ctrl, Het+gRNA 3, Het+gRNAs 2+3, Het+exogenous β3 integrin, KO+gRNA Ctrl and KO+gRNAs 2+3, respectively). **(C)** In Het neurons, the remaining allele of *Itgb3* produces only 50% of the β3 integrin mRNA and protein found in WT neurons, with consequent reduction in network activity. By enhancing expression of the remaining allele, CRISPRa restores β3 integrin levels back to WT values, thus precisely rebalancing network activity. Expression of β3 integrin under the control of an exogenous promoter fails to mimic WT conditions, leading to an un-physiologically high amount of mRNA and protein, thus resulting in hyperactive networks.
